# Inhibition of the HSP90 homodimerization and HSP90-HIF1α interactions by employing small molecules at C-terminal ATP binding site of HSP90

**DOI:** 10.1101/2024.06.02.595921

**Authors:** Jagadeesha Poyya, Chandrashekhar G Joshi

## Abstract

The Heat shock protein HSP 90-alpha (HSP90AA1) is a major chaperone that stabilizes the hypoxia-inducible factor 1α under hypoxic stress and develops solid tumors. Recent studies revealed that in addition to N-terminal ATP binding site, HSP90 has an additional ATP binding site at the C-terminal end. So, disruption of HSP90 and HIF 1 α interaction is an innovative method to control cancer progression. This can be achieved by employing small molecules at C-terminal ATP-binding that do not affect the overall functioning of the HSP90. But there is a lack of structural basis for HIF-1α and HSP90AA1 interactions. This study screened natural products and their derivatives against HSP90AA1 and HIF-1α interaction disruption. Virtual screening was carried out using Glide. We found that compounds with indole rings bind to the C-terminal ATP binding site of HSP90 and prevent its interaction with HIF-1α. Tryptamine derivatives with indole rings showed a greater binding affinity than other molecules. Thus, Tryptamine hydrochloride compounds can be used as a drug repurposing approach to treat hypoxic tumors.

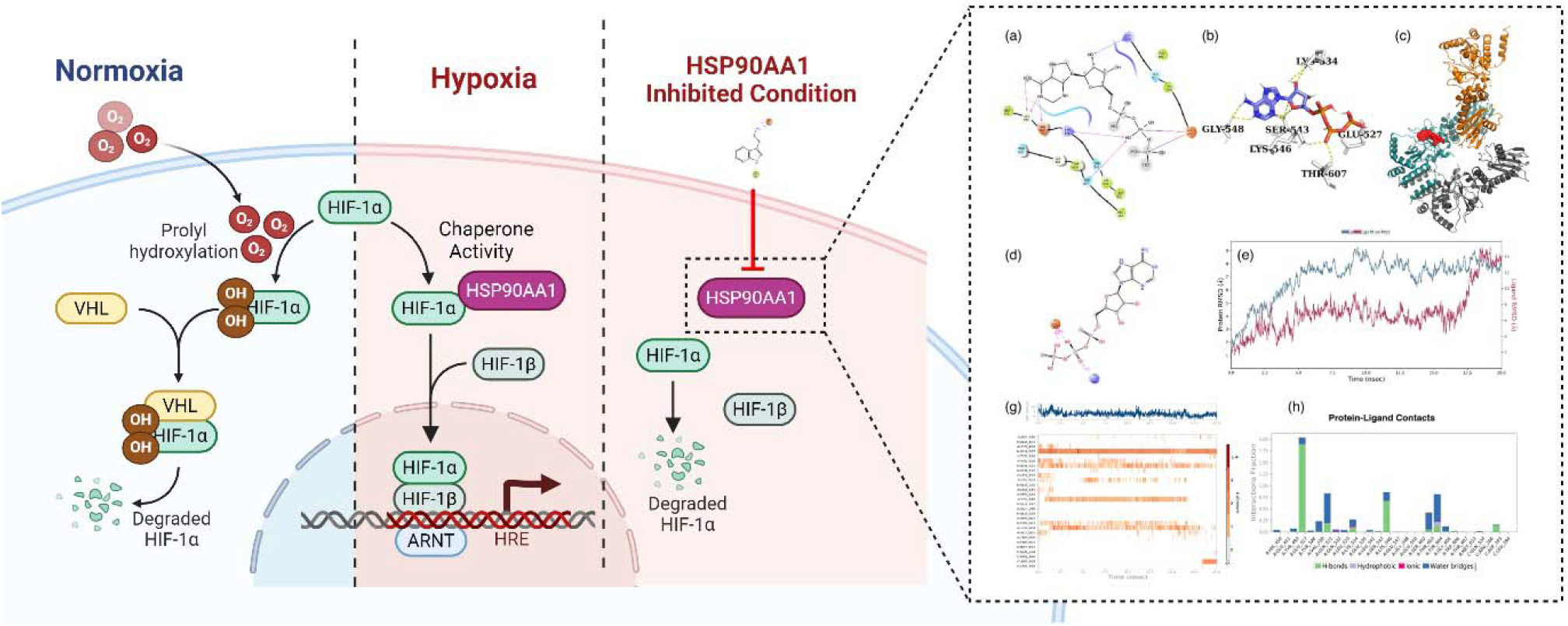

## Introduction

The Heat Shock Protein 90kDa (HSP90) is an ATPase-dependent molecular chaperone expressed when a cell is exposed to proteotoxic stress. Hsp90 controls the refolding or destruction of client proteins by forming a complex with other chaperones such as p23, HSP70, HSP40, and others[1]. The proteins involved in the uncontrolled proliferation and differentiation depend on HSP90 for their folding, stability, refolding, and maturation in tumor cells. Conversely, inhibition of HSP90 induces cellular apoptosis and disruption of multiple signalling pathways, which are involved in the growth and viability of cancer cells. So, new chemicals that can inhibit HSP90 are being discovered to control the cancer menace[2].

HSP90AA1(HSP90) is over-expressed in cancer cells, helping them resist chemotherapy[3]. Studies proved that cancer cell survival rate was correlated with HSP90 expression.[4]. Overexpression of HSP90 was reported in cancers like leukemia and bladder cancer [5,6]. A Tumour biopsy report with a low level of HSP90 is used as a biomarker for successful treatment[7–9]. In osteosarcoma, a chemotherapy agent can induce the expression of HSP90, thereby rendering a chemoresistance in cancer. HSP90 bypassed the PI3K/Akt/mTOR and JNK/P38 pathways, induced apoptosis, and promoted autophagy [10].

In humans, there are five isoforms of Hsp90, including Hsp90AA(inducible) [11]. HSP90 and HSP90AA2 are subtypes Hsp90AA. Out of the two HSP90AA, HSP90 interacts with 376 client proteins and results in proteotoxic stress. This protein contains three structural domains, namely, N-terminal (25kDa), middle (40kDa), and C-terminal domains (12kDa). N-TD performs ATPase activity in these three domains, and MD interacts with various transcription factors, including hypoxia-inducible factor-alpha (HIF-1α).

The HIF-1α is a key transcription factor that optimizes and modifies the cellular function according to the tissue’s oxygen level. The metabolic pathways are reprogrammed in a hypoxic environment through the stabilization and regulation of the degradation of HIF-1 α by the HSP90(Figure 1)[12]. The shift in the glycolysis pathway, angiogenesis, pH homeostasis, metastasis, cell death, and cell survival [13,14] are the events in metabolic reprogramming. Changes in the metabolic pathway make the cancer cells more resistant to chemo and radiation therapy[15]. Consequently, HSP90 inhibitors can be used to overcome the resistance of hypoxic tumors to chemo and radiation therapy. Several studies reported that HSP90 inhibitors could also reduce the HIF-1α expression[16,17]. Hence, HSP90 inhibitors can also be used as HIF-1α negative regulators.

**Figure 1.**
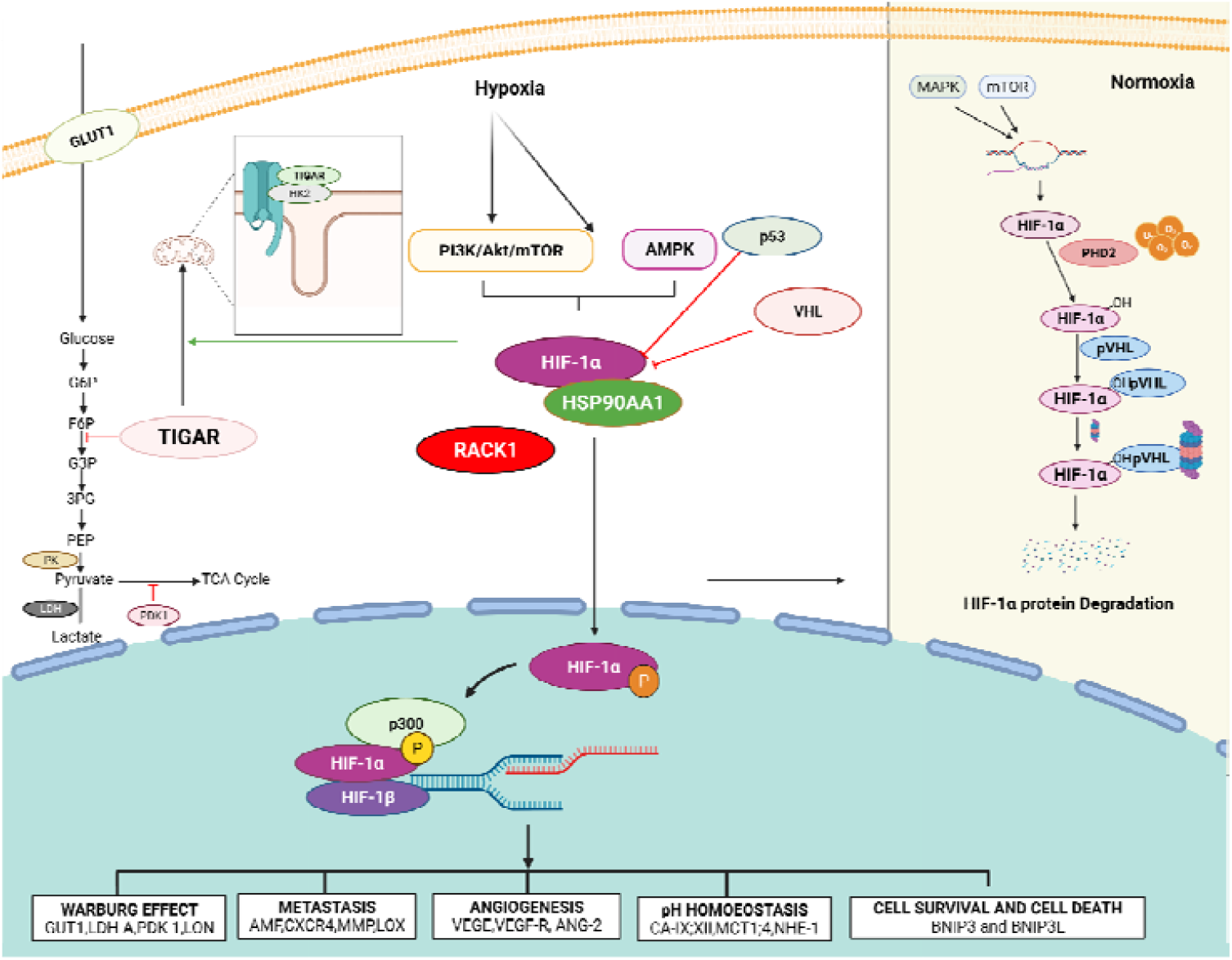
Induction of Hypoxia-inducible factor 1, downstream targets of HIF-1α and its regulation in normoxia. The HIF-1α is an oxygen-sensitive protein, and it undergoes degradation upon hydroxylation by PHD. Impaired expression of the Mitogen-activated Protein Kinase, mammalian target of rapamycin 2, Human-Mouse double minute 2 induces the HIF-1α in cells. Similarly, the mutation in p53 and Von Hippel–Lindau protein prevent the degradation of HIF-1α mediated by proteasome. HSP90AA1 prevents the damage of HIF-1α protein caused by hypoxic stress. RACK1 and HSP90 compete for HIF-1α recruitment. Once HIF-1α becomes stable, it is translocated into the nucleus upon phosphorylation. Translocated protein forms heterodimerization with its β subunits. This heterodimeric complex is recognized by the coactivator p300/CBP and binds to the dimeric complex to form an active complex of HIF-1. The active complex binds the hypoxia response elements (HRE) on the DNA and transcribes the downstream genes. Primary downstream targets of the HIF-1α were genes involving metastasis, angiogenesis, cell survival, cell death, Warburg effect, and pH homeostasis. HIF-1α – hypoxia-inducible factor 1 alpha; PHD2 – Prolyl Hydroxylase 2; vHL-Von Hippel-Lindau; p53-Tumor protein P53; HDM2-Mouse double minute 2 homolog; mTOR-mechanistic target of rapamycin; MAPK-Mitogen-activated protein kinase; HSP90-heat shock protein 90; AMF-autocrine motility factor; C-X-C chemokine receptor type 4; MMP-Matrix metalloproteinase; LOX-Lysyl oxidase; VEGF-Vascular endothelial growth factor; VEGF-R - Vascular endothelial growth factor receptor; ANG-2-Angiopoietin-2; BNIP3-BCL2/adenovirus E1B 19 kDa protein-interacting protein 3; BNIP3L-BNIP3 like protein; GLUT1-Glucose transporter 1; LDHA-A-Lactate dehydrogenase A; PDK1-Pyruvate Dehydrogenase Kinase 1; LON-Lon protease family, CAIX-Carbonic anhydrase IX; CAXII-Carbonic anhydrase XII; MCT1-Monocarboxylate transporter 1, MCT4-Monocarboxylate transporter 4; NHE1-Sodium–hydrogen antiporter 1.

Many compounds have been tested against the N-TD of HSP90. However, during clinical trials, the N-terminal ATP binding site inhibitors showed adverse effects like induction of heat shock response, retinopathy, and gastrointestinal toxicity [18,19]. The limitation of NTD inhibitors promoted researchers to look for a C-terminal ATP binding site. C-TD is responsible for the homodimerization of Hsp90. So, when inhibitors bind to C-TD, dimerization of HSP90 is blocked, preventing it from client protein interactions[20].

Moreover, C-TD can be a selective druggable site as it lacks ATPase activity like N-TD[21]. In addition, C-terminal ATP binding site inhibitors are less harmful compared to N-terminal ATP binding sites. The inhibitory effect was limited to glucocorticoid receptors and not Raf-1, Lck, and C-Src signaling pathways[21]. Therefore, the C-terminal ATP binding site can be an alternative drug target with low side effects. Even though C-TD of HSP90 has fewer side effects, to the best of our knowledge, only a few inhibitors have been tested against this domain [21][22]. In this study, we have screened natural products and their derivatives against the C-terminal ATP binding site of HSP90.

The binding of C-TD inhibitors will allosterically block the binding of ATP at N-terminus. In this study, we hypothesize that employing small molecules at the C-terminal ATP binding site of the HSP90 may reduce the homodimerization of HSP90, HIF-1α-HSP90 interaction, and HIF-1α transactivity.[16, 17].

## Methods

### Protein Preparation

The crystal structure of HSP90 (PDB ID: 3Q6M) was retrieved from Protein Data Bank (http://www.rcsb.org/pdb/home/home.do)[25]. Protein Preparation Wizard tool in Schrodinger with OPLS3 forcefield was used for optimization and minimization of protein structure. This wizard corrects the bond order of amide, hydroxyl, and aromatic groups present in the protein. It also restores the charges and orientations. OPLS3 forcefield minimizes the steric clashes and strains in the protein and the macromolecule’s energy [27].

### Ligand Preparation

Docking was carried out using natural compounds and their derivatives. Totally 67,823 structures were retrieved, and the Lipinski rule of five was applied for screening the large library. Compounds were processed with the LigPrep tool for Glide[27–29] and open babel for AutodockVina, which creates accurate and energy-minimized 3-dimensional structures while applying advanced algorithms to fix the Lewis structure and remove ligand structural errors.

### Active site and Grid generation

Based on an earlier *in-silico* study, we chose the activity of HSP90. There are three binding pockets, namely pck1b, pck2, and pck3[30]. A grid was constructed around the active residues that are reported by Matts et al. based on Novobiocin binding site [31], using the Glide module of the Schrodinger suite’s “Receptor Grid Generation.” The grid was created with an internal dimension of 16×16×16Å (XYZ) and was large enough to contain the protein’s active region, allowing each ligand to search for a possible binding site. In Autodockvina, docking was also used to predict ATP binding. The maximum grid size was created to find a likely ATP binding site. The molecules were docked in Auto Dock Vina and Glide tools to compare the findings.

### Molecular Docking/Virtual Screening

The optimized structure of HSP90 was used to conduct protein-ligand docking experiments. In total, 67,823 molecules were docked against HSP90. The Virtual Screening Workflow docks many chemicals against a single target with three level of precision, such as, HTVS, standard precision, and extra precision by Glide.

### Post Docking MM/GBSA calculations

We used Molecular Mechanics-Generalized Born Surface Area (MM/GBSA) analysis to assess docking efficiency. The Prime MM/GBSA [32,33]uses the MM/GBSA calculation to find the binding free energy (G_*bind*_) of F3 and its derivatives for each ligand and its complex with RdRp.

### Molecular Dynamics Simulation

We employed two alternative tools for Molecular Dynamics (MD) simulation. One of them was D. E. Shaw’s Desmond module using Maestro of Schrödinger [34] suite 2020-1. The Ubuntu environment, which is integrated with NVIDIA T4, was used for Desmond. An MD Simulation study of 20 nanoseconds (ns) was undertaken to investigate the stability of the docked complex with the ligands [35].

The Desmond module of Schrödinger suite 2020-1 was used to investigate the complex in the explicit solvent system with the OPLS2005 force field [36]. For a 10-buffer region, the molecular system was solvated with crystallographic water (TIP3P) molecules under cubic periodic boundary conditions. The system was normalized by introducing Na+ and Cl- as counter ions, which removed the overlapping water molecules. To keep the temperature (300 K) and pressure (1 bar) of the systems constant, an ensemble (NPT) of the Nose–Hoover thermostat [37] and barostat was used. The researchers used a hybrid energy minimization approach that included 1000 steps of steepest descent followed by conjugate gradient algorithms.

We deployed the Gromacs software, version 2018, with the MM/PBSA module[38] to calculate the binding free energy components for Molecular mechanics/Poisson-Boltzmann surface area (MM/PBSA). As described in the previous paper, MM/PBSA calculations of protein-ligand complexes were performed[39].

### Post MD Simulation MM/GBSA analysis

The thermal MM/GBSA.py script from the Prime/Desmond module of the Schrodinger suite 2020-1 was used to perform MM/GBSA analysis after the simulation[40]. For binding free energy calculations of each compounds, every 10th frame from the last 20 ns of simulated trajectories was collected from each MD trajectory, averaging over 50 frames. The Prime MM/GBSA approach employs the rule of additivity, in which total binding free energy (Kcal/mol) is the sum of different energy modules such as coulombic, covalent, hydrogen bond, van der Waals, self-contact, lipophilic, solvation, and ligand and protein π-stackings.

## Results and Discussion

HIF-1α and HSP90 interaction stabilize the HIF-1α during hypoxic stress[41]. During hypoxia, RACK1 and HSP90 compete each other for binding to HIF-1HIF-1α. So, the level of HIF-1α expression decides its binding to HSP90. [42]. Interaction of HSP90 and HIF-1α requires formation of HSP90 homodimerization and active complex[43]. Formation of homodimerization is crucial step in HSP90 active complex formations and it requires binding of ATP at C-terminal ATP binding site. Hence, binding of ATP at C-terminal binding site is check point in HSP90 homodimerization and it can be utilised to develop an inhibitors. Interactions between HIF-1α and HSP90 can break by placing small molecules at the C-terminal ATP binding site[44,45]. Based on this, we designed a virtual screen workflow to screen the HSP90 C-terminal ATP binding site inhibitors to destabilize the HSP90-HIF-1α interactions.

### Molecular Docking/Virtual Screening

HSP90 is required to develop malignancy in tumor cells, and more than 40 oncogenic target proteins have been discovered to date[7,10]. HSP90 inhibitors are also the negative regulators of its client protein, HIF-1α. So, using a single active molecule that could act against both HIF-1α and HSP90 can be a potential drug for cancer treatment.

The crystal structure of the interaction of ATP with the C-terminal ATP binding site of HSP90 has not been studied so far. However, *in-silico* studies established the interaction of the C-terminal ATP binding site of HSP90α with ATP[30,46]. As Novobiocin is a known inhibitor of HSP90, we tried to explore the binding site of ATP on HSP90. Our studies showed that Phosphate and nitrogen of ATP interact with the C-terminal ATP binding site of HSP90 through GLU531 and two Lysine residues (534,546) (Figure 2(a)). The amino acids interacting with ATP and other screened compounds were tabulated in Table 1. The binding energy of screened inhibitors was found to be close to ATP. More details of Novobiocin not shows in the study, as there are many studies were proved its binding site at C-terminal domain of the HSP90.

**Table 1.**
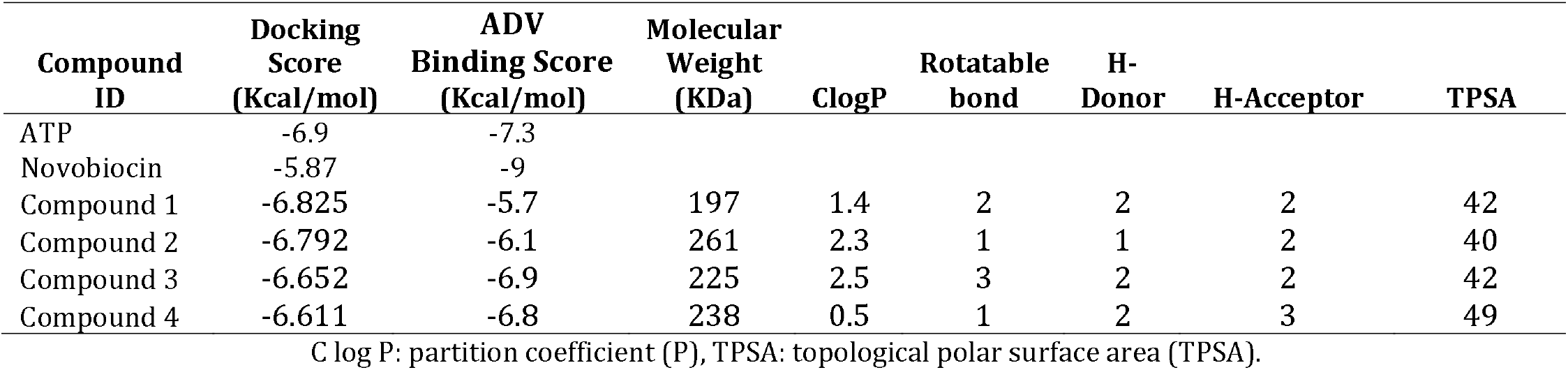
Prioritized compounds that show strong binding interactions with HSP90 and their ADME parameters.

**Figure 2.**
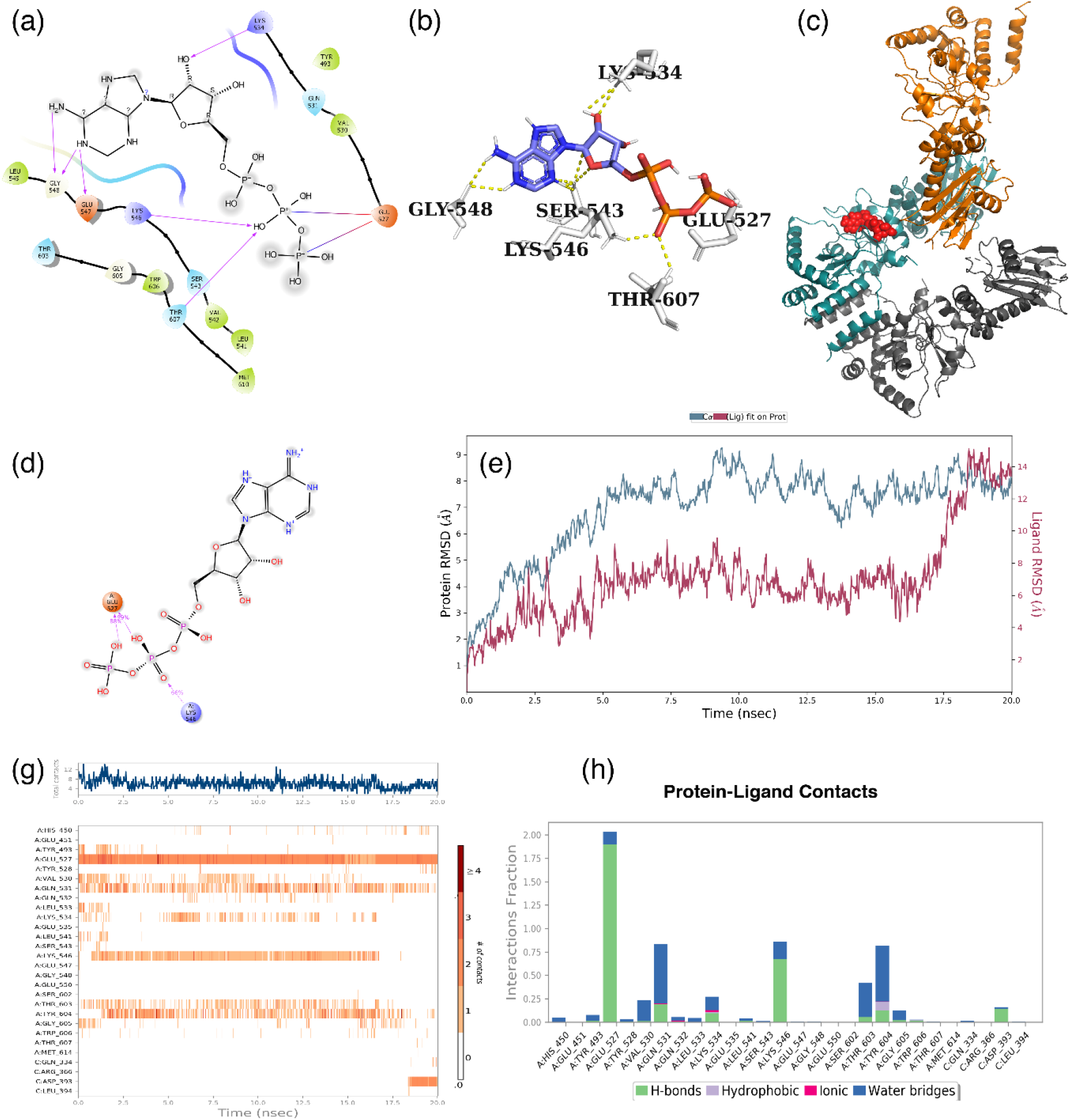
Molecular docking and simulations results of ATP with HSP90. (a) Docked pose of ATP at C-TD of HSP90, (b) ball and stick model representing the interactions of ATP and HSP90. (c) ATP at C-TD binding site of HSP90, where deep teal representing chain A, dark grey showing chain B, chain C showed in orange colour, and red spheres representing ATP. (d) post MD simulation molecular interaction between ATP and HSP90. (e) RMSD of carbon alpha and ligand fit shows stable in their structure throughout the simulations. (f) Protein Ligand Contacts Timeline shows that GLU527, and LYS546 were constantly interacting with ATP during the simulation. (g) Histogram of HSP90 and ATP showing the interactions during the simulation and its interactions fractions.

We have selected indole derivatives as leads out of the screed compounds as they can act on various targets in cancer cells and have shown promising results in the treatment of drug-resistant malignancies. Furthermore, indole-containing drugs including Semaxanib, Sunitinib, Vinorelbine, and Vinblastine have previously been used in clinics to treat a variety of cancers, including drug-resistant ones. As a result, indole derivatives are valuable resources for developing novel lead molecules against drug-resistant cancer.[47]. Recently, it has been shown that inhibitors containing indole groups that bind to C-TD enhanced the anti-proliferative activity against breast and colon cancer cell lines [48,49]. So, we were curious to explore further to find out a potent indole group containing inhibitors against HSP90.

Compound 1 is [2-(1H-indol-3-yl)ethanamine hydrochloride], also known as Tryptamine hydrochloride, which is a proven anti-plasmodial compound [50]. Compound 2 is [2-(5-ethyl-1H-indol-3-yl) ethanamine hydrochloride], and Compound 3 is [2-(5-Propyl-1H-indol-3-YL) ethanamine hydrochloride were surprisingly having similar interaction with HSP90. Compounds 1, 2, and 3 were hydrogen-bonded to GLU527 of HSP90 through an amide linkage. Similarly, the nitro group of indole ring present in these molecules is hydrogen-bonded to TYR604(Figure 3(a), 4(a), and 5(a)). LYS546 was also found to be involved in the interaction with ATP. Detail of hydrogen bonding of selected lead was tabulated in Table 2.

**Table 2.** Hydrogen bonding distance between Compounds with HSP90 and atom involved in the interaction.

| Interactions | Hydrogen Bond Distance in Å |
| --- | --- |
| **ATP** |  |
| A: LYS546:HZ2 - ATP: O12 | 2.207 |
| A: THR607:HN - ATP: O12 | 2.495 |
| ATP:H34 - A: GLU547:O | 2.141 |
| ATP:H34 - A: GLY548:O | 2.404 |
| ATP:H39 - A: GLY548:O | 2.448 |
| **Novobiocin** |  |
| C: LYS632:HN - Novobiocin | 2.301 |
| Novobiocin - C: SER677:O | 2.174 |
| **Compound 1** |  |
| COMPOUND 1:H13 - A: TYR604:O | 1.710 |
| COMPOUND 1:H16 - A: GLU527:O | 2.115 |
| COMPOUND 1:H25 - A: GLU527:OE2 | 1.482 |
| **Compound 2** |  |
| COMPOUND 2:H15 - A: TYR604:O | 1.674 |
| COMPOUND 2:H21 - A: GLU527:O | 2.125 |
| COMPOUND 2:H31 - A: GLU527:OE2 | 1.482 |
| **Compound 3** |  |
| COMPOUND 3:H16 - A: TYR604:O | 1.706 |
| COMPOUND 3:H22 - A: GLU527:O | 2.114 |
| COMPOUND 3:H34 - A: GLU527:OE2 | 1.481 |
| **Compound 4** |  |
| A: TYR493: HH - COMPOUND 4: O11 | 1.942 |
| A: LYS546:HZ1 - COMPOUND 4: O11 | 2.220 |
| COMPOUND 4:H14 - A: GLU527:OE2 | 1.769 |

**Figure 3.**
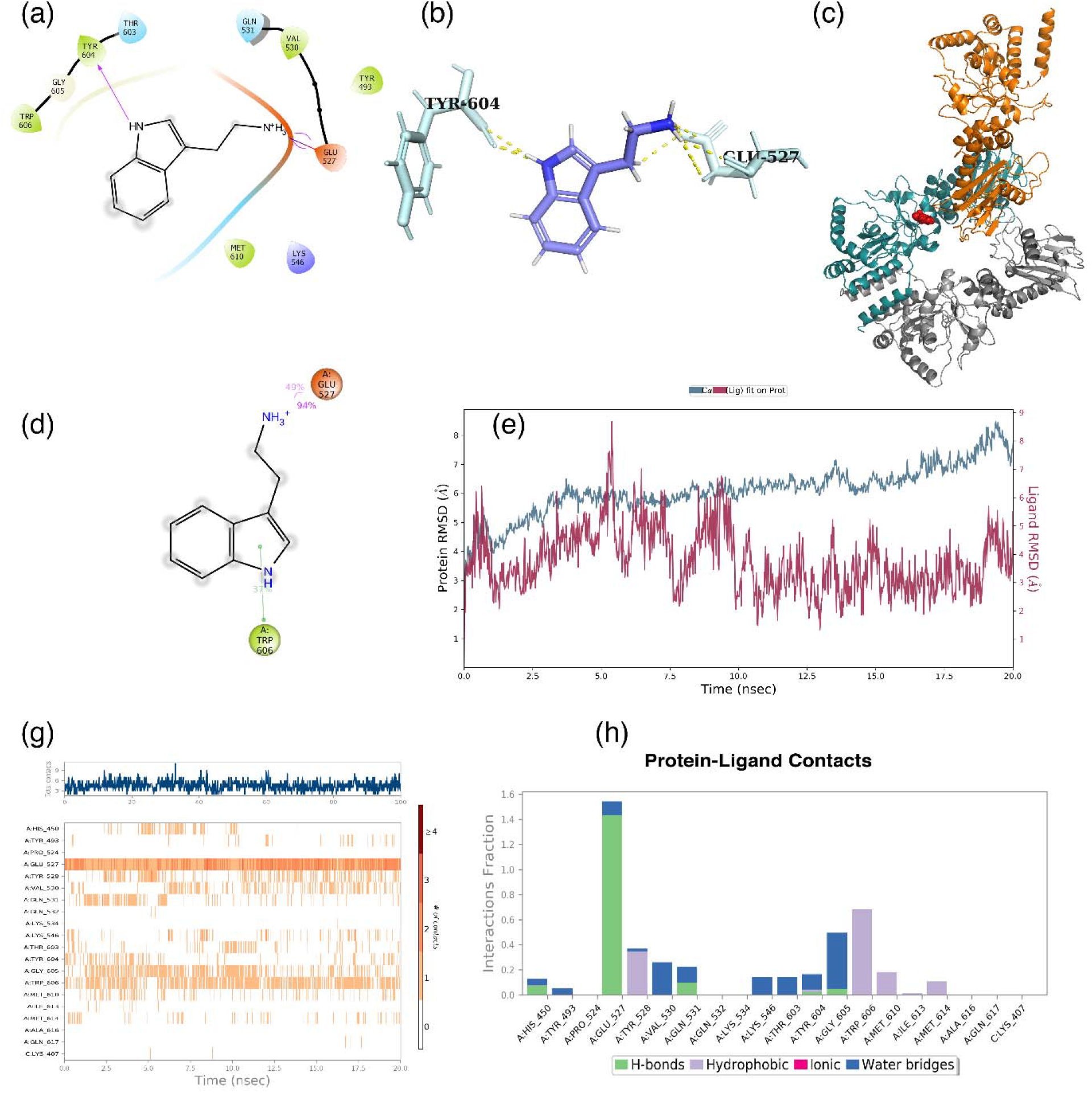
Molecular docking and simulations outcomes of compound 1 with HSP90. (a) Docked pose of compound 1 with HSP90, (b) ball and stick model indicating the interfaces of compound 1 along with HSP90. (c) compound 1 at C-TD binding site of HSP90, where deep teal correspond to chain A, dark grey presenting chain B, chain C showed in orange colour, and red spheres representing compound 1. (d) post MD simulation molecular interface between compound 1 and HSP90. (e) RMSD of carbon alpha and ligand shows the stability in their structure during the course of the simulations. (f) Protein Ligand Contacts Timeline illustrates that GLU527, and TYR606 were continuously interacting with compound1 during the simulation. (g) Histogram of HSP90 and compound 1 showing the interactions during the simulation and its interactions fractions.

Molecular Docking scores were validated by post docking MM/GBSA analysis. This analysis provides clear evidence of the false positive or false negative if any is there in docking studies. The result was tabulated in Table 3, and selected compounds showed more affinity in terms of binding energies. The glide XP score was obtained for each compound, and MM/GBSA complemented each other.

**Table 3.** Post Docking and Post MD simulation MM/GBSA analysis of ligand, protein and its complexes by using Prime module of Schrodinger. Post Docking MM/PBSA was analysed using.

| Compounds | Post-Docking MM/GBSA<br>(Kcal/mol) | Post MD Simulation MM/GBSA<br>(Kcal/mol) | Post MD Simulation MM/PBSA<br>(Kcal/mol) |
| --- | --- | --- | --- |
| ATP | -74.58 | -176.02 | -185.63 |
| Novobiocin | -72.25 | -160.49 | -163.33 |
| Compound 1 | -70.24 | -174.76 | -169.78 |
| Compound 2 | -68.24 | -157.94 | -146.00 |
| Compound 3 | -70.84 | -150.77 | -145.21 |
| Compound 4 | -71.01 | -131.42 | -130.74 |

*g\_mmpbsa* module of Gromacs.

Further, we conducted blind docking using AutoDock Vina to compare the binding site. The binding score and interacting amino acids were the same as glide results. Hence, these studies showed that selected compounds could efficiently bind to the HSP90 C-terminal ATP binding site. Further, only three compounds were selected for the MD simulation and along with ATP.

### Molecular Dynamics Simulations

We carried out MD simulations to determine the stability of protein, ligand, and their interactions during certain physiological conditions. Based on the glide XP score and post docking MM/GBSA, only three ligand interactions with HSP90 were studied in MD simulations. MD simulations of small molecule and HSP90 were carried out for about 20000ps. Interestingly, RMSD, and RMSF of all the four compounds, and ATP were stable during the simulations.

The interactions fractions index of ATP and HSP90 interactions showed that GLU527 contributed to hydrogen bonding and water bridges. The stability graph showed two interactions, where the second phosphate group of ATP contributed about 99%, and the third was about 88% in hydrogen bonding (Figure 3(d)). These two interactions between GLU527 and ATP were stable throughout the simulations (Figure 3(g)). Another residue, LYS546, donated the hydrogen bond to the phosphate group of ATP, and this interaction remained only for 80% of the simulations. A water bridge was also found between LYS546 of HSP90 and ATP. These results infer that GLU527 and LYS546 are the ATP interacting residues of HSP90, as was confirmed by per residue interactions fractions (Figure 3(h)).

Docking studies and post docking MM/GBSA studies revealed that Compound 1 was potent enough to bind to the C-terminal ATP binding site of HSP90. The interaction observed during docking was the same except for TYR604. Instead of TYR604, TRP606 was in contact with compound 1 with pi-pi stacking. Compound 1 had two salt bridges with GLU527, and this bond was very stable throughout the simulation.

During the simulation, interactions between ARG400, GLU401, LYS407, and TYR528 of HSP90 with compound 2 were stable (Figure 4(g)). Per residue binding fractions showed that the hydrogen bond donated by ARG400 and LYS407 to the phosphate group of compound 2 contributed about 91% of its interactions. LYS407 interactions with compound 2 were stable throughout the simulations (Figure 4(d)).

**Figure 4.**
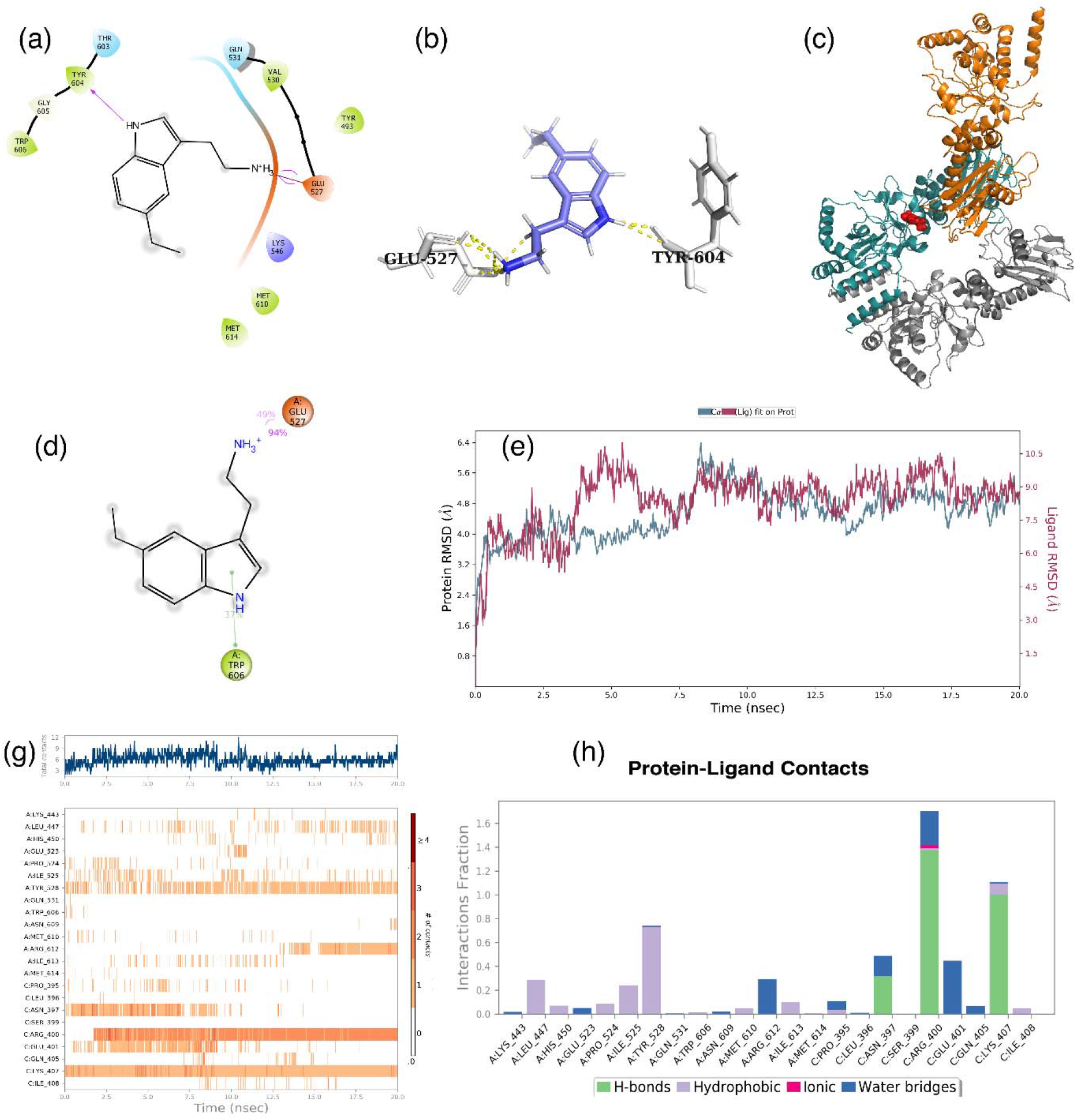
Molecular docking and simulations figures of compound 2 with HSP90. (a) two-dimensional representation of interactions between compound 2 with HSP90, (b) binding site residues of HSP90 were interacting with compound 2 and it is represented in ball and stick model. (c) carton model of the compound 2 bound HSP90 showing binding of compound 2 at C-TD, where deep teal correspond to chain A, dark grey representing chain B, chain C showed in orange colour, and red spheres representing compound 2. (d) schematic representation of molecular interactions between compound 2 and HSP90 during post MD simulation. (e) RMSD graph showing the stable protein structure and ligand throughout the progression of the simulations. (f) Timeline of ligand-protein interactions shows that GLU527, and LYS546 were unceasingly interacting with compound1 in the simulation. (g) Effectiveness of each interactions in term of interaction fraction between compound 1 and HSP90 represented in a histogram.

In addition to this, ARG400 of HSP90 donates one more hydrogen bond, which is less stable during the simulation and contributes to only 45% of interactions with compound 2. TYR528 in chain A of HSP90 was involved in pi-pi stacking with compound 2, and this bond was less stable. A water bridge was found between GLU401 and compound 2, contributing only 33% of HSP90 and compound 2 interactions. (Figure 4(h))

Compound 3 had interactions with GLU527 as confirmed during MD simulation. GLU527 showed contact with compound 3 through the hydrogen bond, ionic, and water bridges during the MD simulation analysis (Figure 5(h)). GLU527 formed two salt bridges between the amino group of compound 3((Figure 5(d))). These interactions were stable during the simulation (Figure 5(g)). The second important interaction was GLN531 which accepts one hydrogen bond from the amino group of compound 2. Finally, TRP606 was formed pi-pi stacking with compound 3. The data obtained from the MD simulation confirmed the stable interaction between compounds 1,2, and 3 with C-terminal ATP binding site residues of HSP90((Figure 3(c), 4(c), and 5(c)).

**Figure 5.**
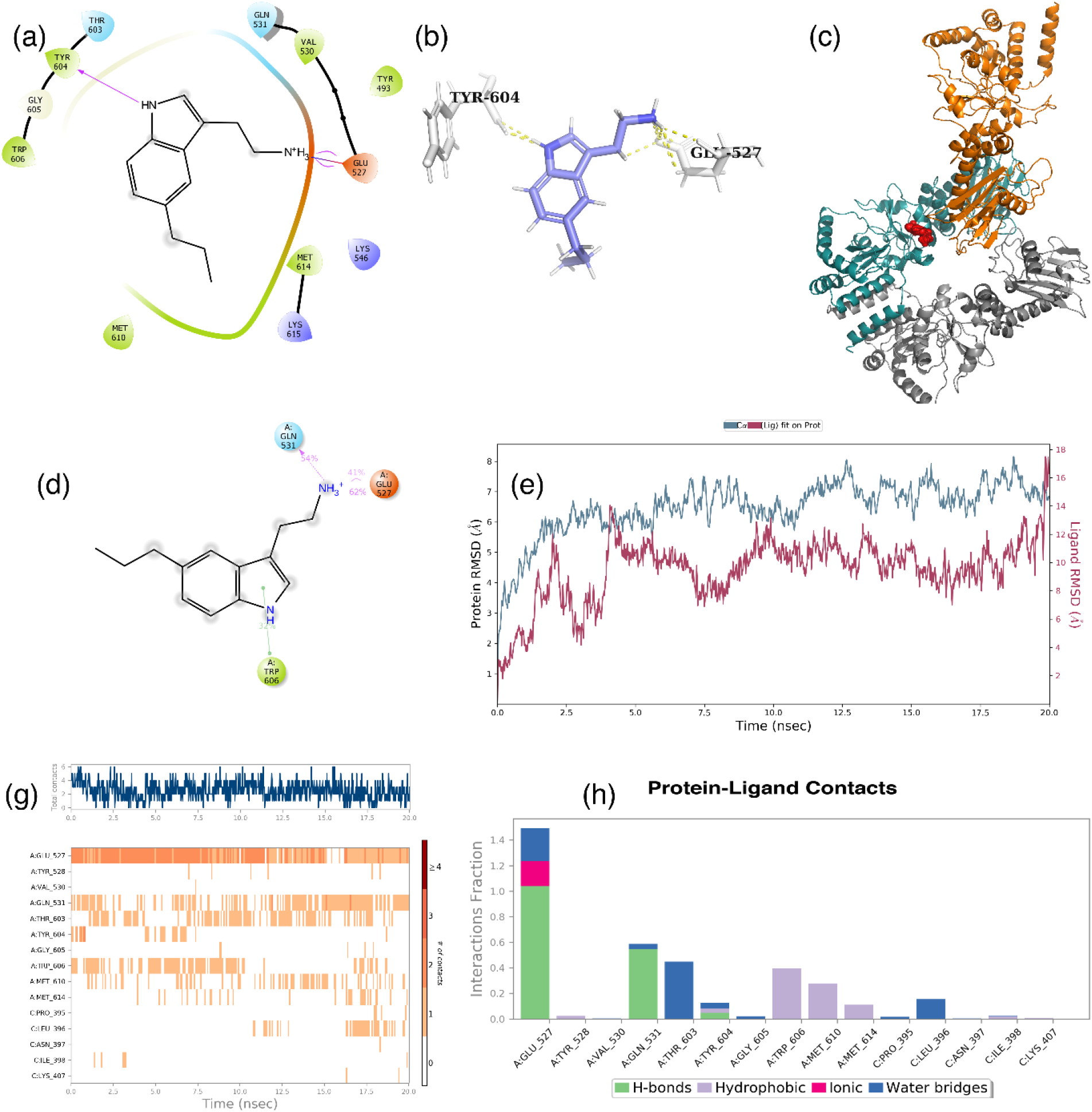
Molecular docking and simulations data of compound 3 with HSP90. (a) Pose view of docked compound 3 and HSP90, (b) interacting residues where represented in ball and stick model. (c) Carton model of the HSP90 showing binding of compound 3 at C-TD, where deep teal correspond to chain A, dark grey representing chain B, chain C showed in orange colour, and red spheres representing compound 3. (d) Depiction of post MD simulation molecular interface between compound 3. (e) Deviation in the alpha carbon atom of the compound and protein infer that protein structure and ligand were experience less conformation changes during the simulation. (f) Timeline of ligand-protein interactions shows that GLU527, GLN531, and TRP604 were perpetually interrelating with compound1 during the simulation. (g) Efficacy of individual contacts in term of interaction fraction among compound 1 and HSP90 characterized in a histogram.

### Binding energy estimation of MD trajectories using MM/GBSA and MM/PBSA

The average binding energies with standard deviation were listed in Table 3 after post-simulation MM/GBSA was computed for every 10th frame, comprising 50 conformations of each simulated complex. Because it considers protein flexibility, entropy, solvation, and polarizability, which are usually unaccounted for in several docking protocols, MM/GBSA is a very popular and rigorous method for post-simulation binding free energy prediction. As a result, it is reliable than that of most scoring functions used during molecular docking. One of the most significant tasks in biomolecular studies is to calculate the free energy of binding precisely because it is responsible for driving all molecular processes such as chemical reactions, molecular recognition, association, and protein folding.

The *g_mmpbsa* script was used to do the binding free energy computations. New trajectories comprised of the final 10ns trajectories of the output trajectory were built using the gmxtrjconv module, with frames generated every 200ps, for the Molecular mechanics/Poisson-Boltzmann surface area (MMPBSA) calculations. The energies for each compounds were tabulated in Table 3.

Both post MD simulation MM/GBSA and MM/PBSA showed that the compound 1 (−185.63 Kcal/mol) was closer to the ATP and Novobiocin in terms of binding free energies (Table 3). Molecular interactions during docking, MD simulation and binding free energies of selected lead proposed a new C-terminal ATP binding inhibitors. Low binding energies of compound 4 had made us drop it during MD simulation discussion.

Previous studies reported that HIF-1α interacts with Hsp90 through the bHLH-PAS domain under normoxia. However, the activation of HIF-1α in hypoxia requires the presence of an active Hsp90[51] which inhibits the ubiquitination of former and proteasomal degradation. It has been recently shown that HIF-1α expression can be reversed using HSP90 siRNA, leading to the apoptosis and death of cells[52].

The C-terminal inhibitors of HSP90 can cause proteasomal degradation of HIF-1α during hypoxia. Consequently, downstream target genes of HIF-1α, especially GLUT1, Hexokinase 2, and TIGAR expression, were suppressed[17][53]. Increased glucose uptake and GLUT1 expression are linked to the worsening survival chances of cancer patients[54]. C-TD inhibitors have been shown to downregulate HK2 in colorectal, breast, and prostate cancer cells under the influence of HIF-1α. So, we hypothesize that the HSP90 inhibitors can also act against HIF-1α and prevent the progression of cancer.

In summary, the different sets of compounds were found to bind at pocket3 of the C-terminal domain of HSP90. These compounds were maybe promising inhibitors that could inhibit the HSP90-HIF-1α interactions. Obstruction in this interaction may cause structural changes in HIF-1α under the hypoxic stress, resulting in its degradation. These compounds obey Lipinski’s rule of five and are less toxic to non-cancerous cells. Our study showed the possible role of these molecules in preventing the progression of cancer by blocking the HSP90 homodimerization and HSP90-HIF1α interactions.

## Acknowledgement

Authors acknowledge the Google Cloud Education Programs Team for providing Google Cloud Research Credits (EDU Credit wilsonjessica 209474439). And, Department of Studies in Biochemistry, Mangalore University, for the facility and support.

